# Regional oceanographic features and hydrothermal activity influence protist diversity and biogeography in the Okinawa Trough

**DOI:** 10.1101/714816

**Authors:** Margaret Mars Brisbin, Asa E. Conover, Satoshi Mitarai

## Abstract

Microbial eukaryotes (protists) contribute substantially to ecological functioning in marine ecosystems, but factors shaping protist diversity, such as dispersal barriers and environmental selection, remain difficult to parse. Deep-sea water masses, which form geographic barriers, and hydrothermal vents, which represent isolated productivity hotspots, are ideal opportunities for studying the effects of dispersal barriers and environmental selection on protist communities. The Okinawa Trough, a deep, back-arc spreading basin, contains distinct water masses in the bottom waters of northern and southern regions and at least twenty-five active hydrothermal vents. In this study, we used metabarcoding to characterize protist communities from fourteen stations spanning the length of the Okinawa Trough, including three hydrothermal vent sites. Significant differences in community structure reflecting regional oceanography and water mass composition were present, indicating the importance of geographic factors in shaping protist communities. Protist communities in bottom waters affected by hydrothermal activity were significantly different from communities in other bottom waters, suggesting that environmental factors can be especially important in shaping community composition under specific conditions. Amplicon sequence variants that were enriched in hydrothermally influenced bottom waters largely derived from cosmopolitan protists that were present, but rare, in other near-bottom samples, thus highlighting the importance of the rare biosphere.

## Introduction

Microbial unicellular eukaryotes (protists) are important contributors to all marine ecosystems, from the sunlit surface waters [1] to the deep, dark bathypelagic [2]. Extensive phylogenetic and functional diversity among protists—especially in extreme environments like the deep sea and hydrothermal vents [3]—has been revealed by high-throughput sequencing with environmental samples, but factors influencing protist community structure remain difficult to parse [4, 5]. Physicochemical factors, such as temperature, light and nutrient availability, have historically been regarded as having the strongest influence on microbial community structure, but geographic restrictions— between ocean basins [4] and water masses [5, 6]—and biological interactions [7] can also contribute substantially to shaping microbial community structure. Furthermore, physicochemical, biological, and geographical factors can vary in their relative influences on community structure at different depths and in different oceanic regions [8].

The Okinawa Trough (OT), a back-arc spreading basin (> 2000 m) within the East China Sea, is an ideal environment to investigate factors shaping protist community structure. The Kuroshio Current, the western boundary current of the North Pacific subtropical gyre, enters the OT to the east of Taiwan, and transports warm, high-salinity water northward [9]. The bottom water in the southern OT is formed from Kuroshio Intermediate water (KIW), which is 55% high-salinity South China Sea Intermediate Water (SCSIW) and 45% low-salinity North Pacific Intermediate Water (NPIW), flowing over the sill (775 m) east of Taiwan and into the southern OT [10]. Additional surface water and intermediate water are entrained into the OT through the Kerama Gap, which, with a sill depth of 1100 m, is the deepest channel connecting the OT and the Philippine Sea. NPIW consistently enters the deep regions of the OT through the Kerama Gap [11], increasing the contribution of NPIW to bottom waters in the northern OT to 65% [10]. However, surface flow in the Kerama Gap is periodically reversed, likely due to the arrival of mesoscale eddies from the interior of the North Pacific [11]. The topography and current structure in the OT lead to varying environmental conditions in the northern and southern trough, as well as in the Kerama Gap. Most notably, water mass compositions are distinct in the bottom waters of the southern (45% NPIW, 55% SCSIW) and northern OT (65% NPIW, 35% SCSIW) and the Kerama Gap (75% NPIW, 25% SCSIW) [10], thus facilitating study of how water mass composition impacts on protist community structure.

As a back-arc spreading basin, there is extensive hydrothermal activity throughout the Okinawa Trough [12]. Twenty-five active OT hydrothermal vent sites are listed in The InterRidge Database v3.4 [13], but this figure likely remains an underestimate as new vents continue to be discovered [14, 15]. The hydrothermal vents in the OT are distinct from those found at mid-ocean ridges, primarily due to thick layers of terrigenous sediments overlaying vent sites. Compared to sediment-starved vent systems, OT vent fluids are typically low pH with high concentrations of CO_2_, NH_4_^+^, boron, iodine, potassium, lithium [16] and methane [17]. Unique physicochemical conditions associated with different vent systems—such as those associated with OT vents—give rise to heterogeneous biological communities [18], warranting careful study of diverse hydrothermal systems. In addition, organic matter and inorganic nutrients resuspended by vents in the OT may locally increase microbial production in bottom waters [19]. However, vent fields in the Okinawa Trough have not been studied nearly as extensively as hydrothermal vents in other oceanic regions [12].

In this study, we present the first comprehensive survey of protist community structure in the Okinawa Trough and Kuroshio Current. We analyzed protist community composition in replicate samples collected from the sea surface, subsurface chlorophyll maximum, middle water column, and bottom waters at sites spanning the entire length of the Okinawa Trough and within the Kerama Gap. In addition, bottom waters at three of our sampling sites were influenced by nearby hydrothermal vents, as evidenced by increased turbidity, CDOM (colored dissolved organic matter), NH_4_, and total carbon concentrations—conditions typical of hydrothermal vent plumes in the OT [16, 20]. The main objectives of the study were to: (i) assess broad patterns in protist diversity in the Okinawa Trough and Kuroshio Current, (ii) determine the extent to which OT protist diversity is affected by regional ocean circulation and water mass composition, and (iii) investigate the influence of hydrothermal activity on deep-sea protist communities in the OT.

## Materials and Methods

### Sampling locations

Water samples were collected from 14 sites in the Okinawa Trough and Kerama Gap during the Japan Agency for Marine-Earth Science and Technology (JAMSTEC) MR17-03C cruise from May 29 to June 13, 2017 (Figure 1). Stations 8, 9, 10, and 11 are located in the Southern Okinawa Trough (SOT); stations 3, 4, 2, 5 make up a transect of the Kerama Gap (KG) from east to west; stations 12, 13, 14, 15, 17, and 18 are in the Northern Okinawa Trough (NOT) stations. Five stations are located near known active hydrothermal vent sites; station 10 is at the Hatoma Knoll [16], station 2 is at the ANA vent site of the Daisan-Kume Knoll [14], station 11 is near the Dai-Yon Yonaguni Knoll [21], station 5 is near the Higa vent site [14], and station 12 is near the Iheya North vent field [22]. Physicochemical data collected by in-situ sensors and chemical analysis of water samples were used to discern whether or not samples were influenced by hydrothermal vent plumes. Vertical profiles from stations 2, 10, and 11 showed near-bottom turbidity and CDOM maxima (Supplemental Figure 1A), which is typical for OT vent plumes [16, 20]. In addition, nutrient analysis showed elevated NH_4_ concentrations in bottom water samples from stations 2 and 10, providing further evidence of hydrothermal influence [16, 20] (Supplemental Figure 1B). Total carbon was only measured for samples from stations 2, 10, 14, and 15; consistent with hydrothermal influence [16, 20], stations 2 and 10 exhibit near bottom peaks in carbon concentrations (Supplemental Figure 1C). Stations 2, 10 and 11 are, therefore, included in analyses aimed at determining how hydrothermal activity influences protist community structure in the OT, while the other sampling stations near vent sites are not.

**Figure 1.**
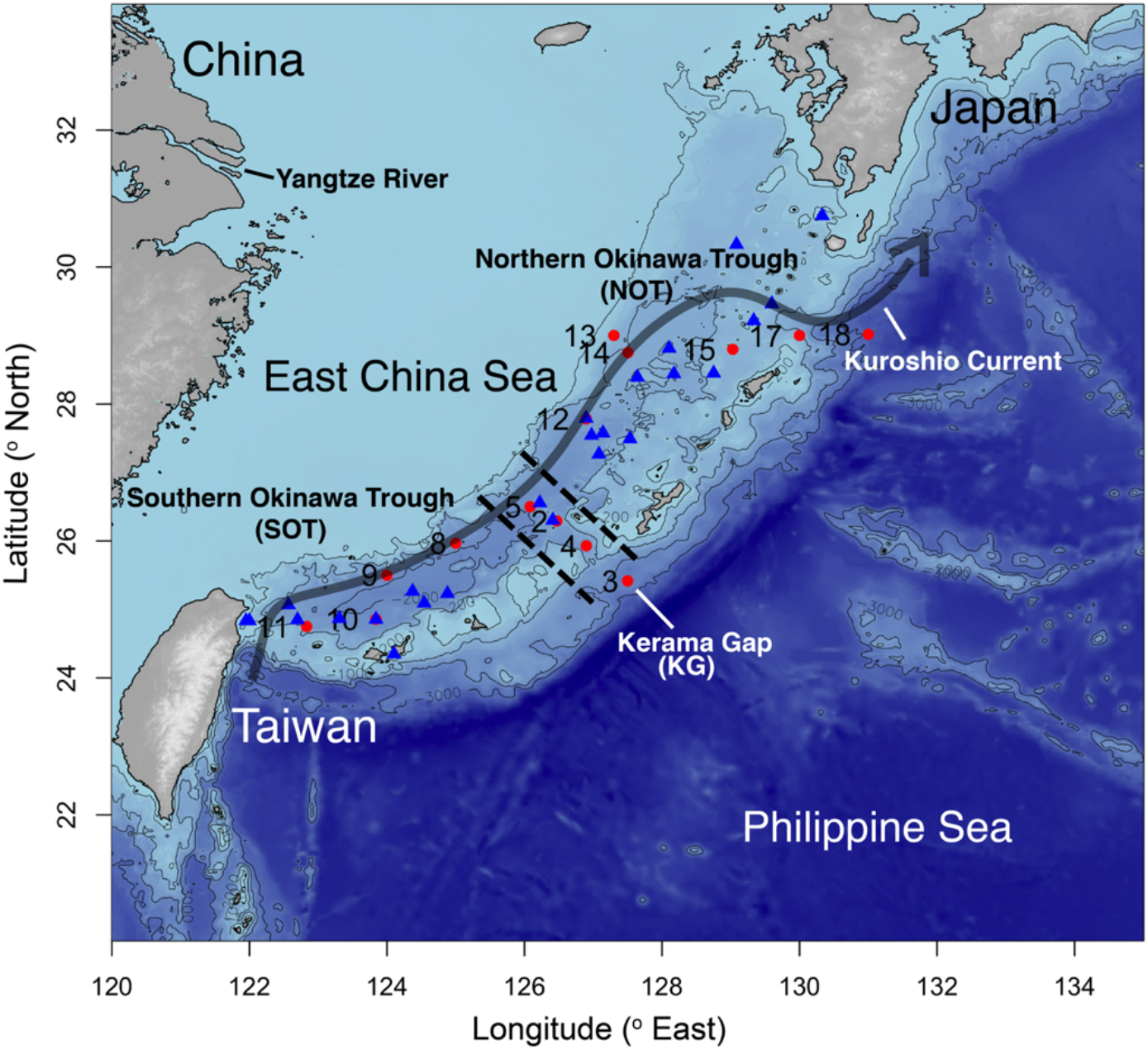
Sampling locations in the Okinawa Trough. Numbered red circles denote sampling stations where water was collected during the Japan Agency for Marine-Earth Science and Technology (JAMSTEC) MR17-03C cruise in May and June 2017. Blue triangles indicate active hydrothermal sites according to the InterRidge Vents Database v3.4 (https://vents-data.interridge.org/ventfields-osm-map, accessed 05/08/2019). Bathymetry data was accessed and plotted through the function getNOAA.bathy in R package marmap. The 200, 1000, 2000, and 3000 m isobaths are plotted and labeled. Station 2 is at the ANA site of the Daisan-Kume Knoll, station 10 is at the Hatoma Knoll, station 11 is near the Dai-Yon Yonaguni Knoll, station 5 is near the Higa vent site, and station 12 is near the Iheya North vent field.

### Sample collection

A Niskin rosette with 30 10-liter bottles and fitted with a conductivity-temperature-depth (CTD) probe (SBE 911plus, Sea-Bird Scientific, Bellevue, WA) was deployed at each cruise station to collect water from the subsurface chlorophyll maximum (SCM, 50–100 m), the middle water column (mid, 700–1500 m), and approximately 10 m above the seafloor (bottom, 772–2957 m) (Supplemental Table 1). Surface seawater was collected by bucket alongside the research vessel. Two replicates of 4.5 liters (surface) from separate bucket casts or 5 liters from separate Niskin bottles (SCM, mid, bottom) were sequentially filtered under a gentle vacuum through 10.0-μm and 0.2-μm pore-size polytetrafluoroethylene (PTFE) filters (Millipore, Burlington, MA). Filters were flash-frozen in liquid nitrogen and stored at −80°C. Temperature, salinity, dissolved oxygen, fluorescence, and turbidity profiles were recorded by CTD probe at each station. CDOM and chemical analyses for nitrate, nitrite, NH_4_, phosphate, silicate and total carbon were performed by Marine Works Japan Ocean Chemistry Analysis Section onboard with water collected by Niskin bottle.

### DNA extraction and sequencing library preparation

DNA was extracted from PTFE filters (n = 224, two replicates of two filter pore-sizes at four depths from 14 stations) following manufacturer’s protocols for the DNeasy PowerWater Kit (Qiagen, Hilden, Germany) including the optional heating step for 10 min at 65°C to fully lyse cells. Sequencing libraries were prepared following the Illumina 16S Metagenomic Sequencing Library Preparation manual, but with universal eukaryotic primers for the V4 region of the eukaryotic 18S rRNA gene (F: CCAGCASCYGCGGTAATTCC [23], R: ACTTTCGTTCTTGATYR [24]) and 58°C annealing temperature in the initial PCR. Amplicon libraries were sequenced by the Okinawa Institute of Science and Technology (OIST) DNA Sequencing Section on the Illumina MiSeq platform with 2×300-bp v3 chemistry. Amplification and sequencing were successful for 211 samples and at least one replicate succeeded for each sample type.

### Amplicon sequence analysis

Sequence data from each of four flow-cells were denoised separately using the Divisive Amplicon Denoising Algorithm [25] through the DADA2 plug-in for QIIME 2 [26]. We analyzed amplicon sequence variants (ASVs) to maximize the amount of diversity included in the study [27] and detect changes in community composition that may not be apparent if analyzed at higher taxonomic levels. Denoised ASV tables were merged before taxonomy was assigned to ASVs with a naive Bayes classifier trained on the Protist Ribosomal Reference (PR2) database [28] using the QIIME 2 feature-classifier plug-in [29]. We imported results into the R statistical environment [30] for further processing with the R package phyloseq [31]. Sequences were initially filtered to remove those that were not assigned taxonomy at the Kingdom level—likely data artifacts—and all metazoan sequences. We did not apply additional prevalence or minimum abundance filtering since samples were from varied locations and depths, but the DADA2 algorithm discards singletons.

### Statistical analyses

To test whether community composition varied significantly by depth, filter-size, region or presence or absence of hydrothermal influence in bottom water samples, we computed the Aitchison distance between samples, which minimizes compositional bias inherent in metabarcoding data [32], and performed Permutational Multivariate Analyses of Variance (PERMANOVA, 999 permutations) with the adonis2 function from the R package vegan [33]. We further used pairwise PERMANOVA (999 permutations) to test which regions differed in community composition from each other. Because stations 3 and 18 were much deeper than other stations (Supplemental Table 1), we used midwater samples from these stations in the bottom-water analyses and excluded these stations from the mid-water analyses. This is further justified because the mid-water samples at these stations were from the salinity layer contributing to bottom waters at the other stations (Supplemental Figure 2). To determine if environmental variables contributed to changes in community composition, we performed Redundancy Analysis (RDA) and applied ANOVA to RDA results to test whether RDA models were statistically significant and which environmental variables significantly contributed to variation in community composition [34]. Before performing RDA, we tested for collinearity of environmental variables by computing Pearson correlation coefficients between environmental variables in each depth layer (Supplemental Figure 3). If the absolute value of correlation coefficients was greater than 0.8, we only included one of the correlated variables in the RDA for that depth [35]. We further applied variance partitioning on variables that significantly contributed to variation in community composition to determine the percent of variation attributable to each variable [36]. Lastly, we used the DESeq2 Bioconductor package [37] to determine which ASVs were differentially abundant in bottom water from sites influenced by hydrothermal activity compared to bottom water at other sites. The data and code necessary to reproduce the statistical analyses are available on github (https://github.com/maggimars/OkinawaTroughProtists), including an interactive online document: https://maggimars.github.io/OkinawaTroughProtists/OTprotists.html.

## Results

### Protist diversity in the Okinawa Trough

Overall, 31.5 million sequencing reads were generated for this study, with 34,631–421,992 sequencing reads per sample (mean = 144,604). All sequences are available from the NCBI Sequencing Read Archive with accession PRJNA546472. Following denoising, 16.8 million sequences remained, with 1,724–215,842 sequences per sample (mean = 77,342). A total of 22,656 unique ASVs were identified in our dataset, with 49–1,906 observed ASVs per sample (mean = 730). We used pairwise Wilcox tests on observed richness and Shannon indices to determine whether alpha diversity differed between depths or was influenced by hydrothermal activity. The SCM had significantly higher richness than surface, mid, and bottom waters (*p* < 0.001, Supplemental Figure 4A). Shannon indices for SCM and surface samples were not significantly different but were both significantly higher than for mid and bottom waters (*p* < 0.001, Supplemental Figure 4B). Hydrothermally influenced bottom waters at stations 11, 10, and 2 did not have significantly different observed richness or Shannon indices from other bottom water samples (Supplemental Figure 4C–D).

Samples clustered by depth in a Principal Coordinate Analysis (PCoA) of Aitchison distance between samples, with SCM and surface samples clustering separately from mid and bottom water samples on the primary axis (Figure 2). SCM and surface samples further separated on the secondary axis, but mid and bottom water samples clustered together. PERMANOVA results were significant at *p* < 0.05 when performed by depth layer on all samples. When samples from each depth were analyzed separately, samples clustered by filter pore-size on the primary axis for surface and SCM samples, and on the secondary axis for mid and bottom water samples (Supplemental Figure 5A–D), but PERMANOVA results by filter pore-size were not statistically significant.

**Figure 2.**
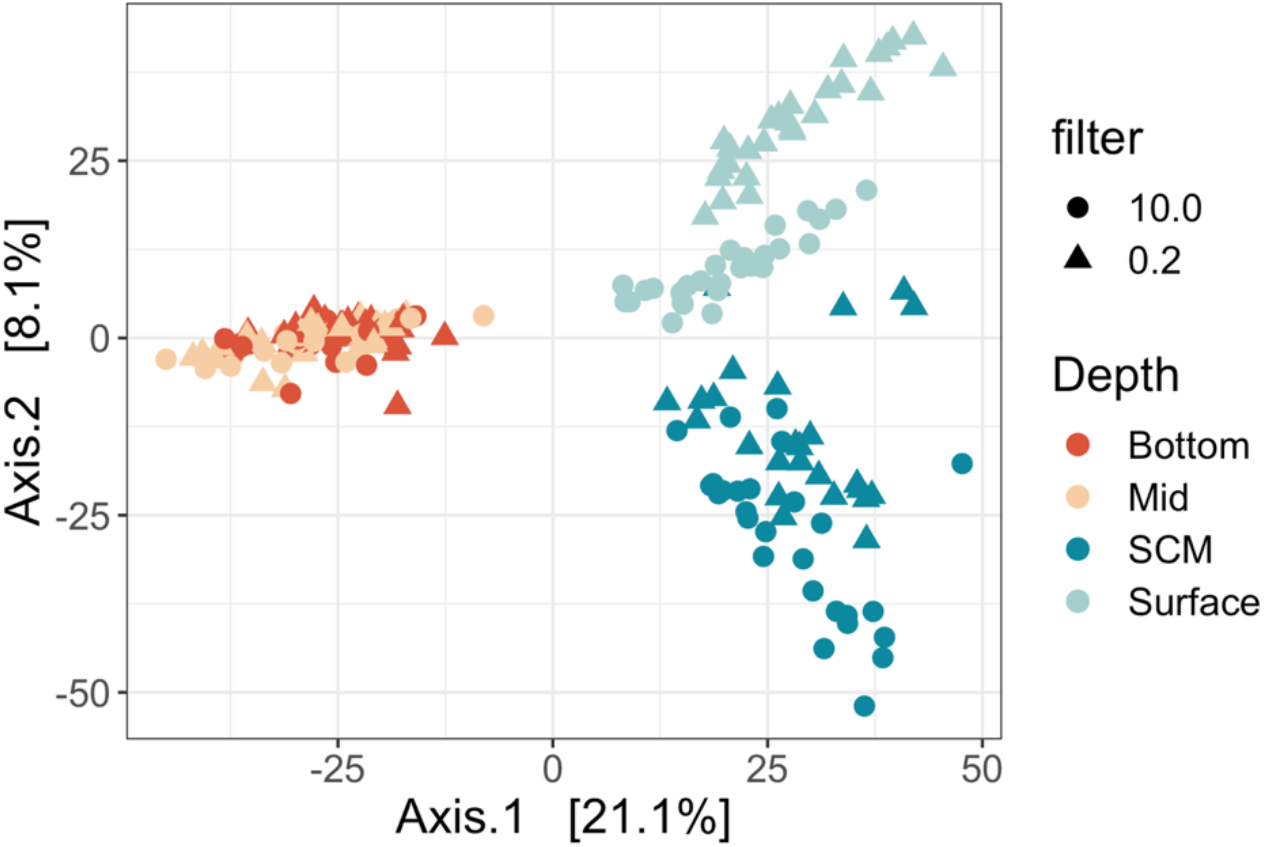
Principal coordinates analysis of Aitchison distances between size-fractionated protist communities from four depths in the Okinawa Trough. Point color indicates the depth layer from which samples were collected and point shape reflects the filter pore-size used to collect samples in μm. Samples cluster in three main groups by depth: surface, subsurface chlorophyll maximum (SCM), and mid/bottom waters. Community composition was significantly different by depth layer (PERMANOVA, 999 permutations, *p* < 0.05).

Surface samples were dominated by Dinoflagellata, Haptophyta, and Radiolaria in the larger size-fraction and Dinoflagellata, Haptophyta, Ochrophyta and MAST groups in the smaller size-fraction (Figure 3). In the SCM, Dinoflagellata, Haptophyta, Ochrophyta and Radiolaria were most abundant in the larger fraction and Chlorophyta, Dinoflagellata, Haptophyta, Ochrophyta, Radiolaria, and MAST groups were most abundant in the smaller size-fraction. The mid and bottom water samples were dominated by Dinoflagellata and Radiolaria in both size-fractions. Interestingly, bottom water samples from hydrothermally influenced stations (2, 10, 11) included noticeably higher proportions of Ciliophora, Ochrophyta, and MAST groups.

**Figure 3.**
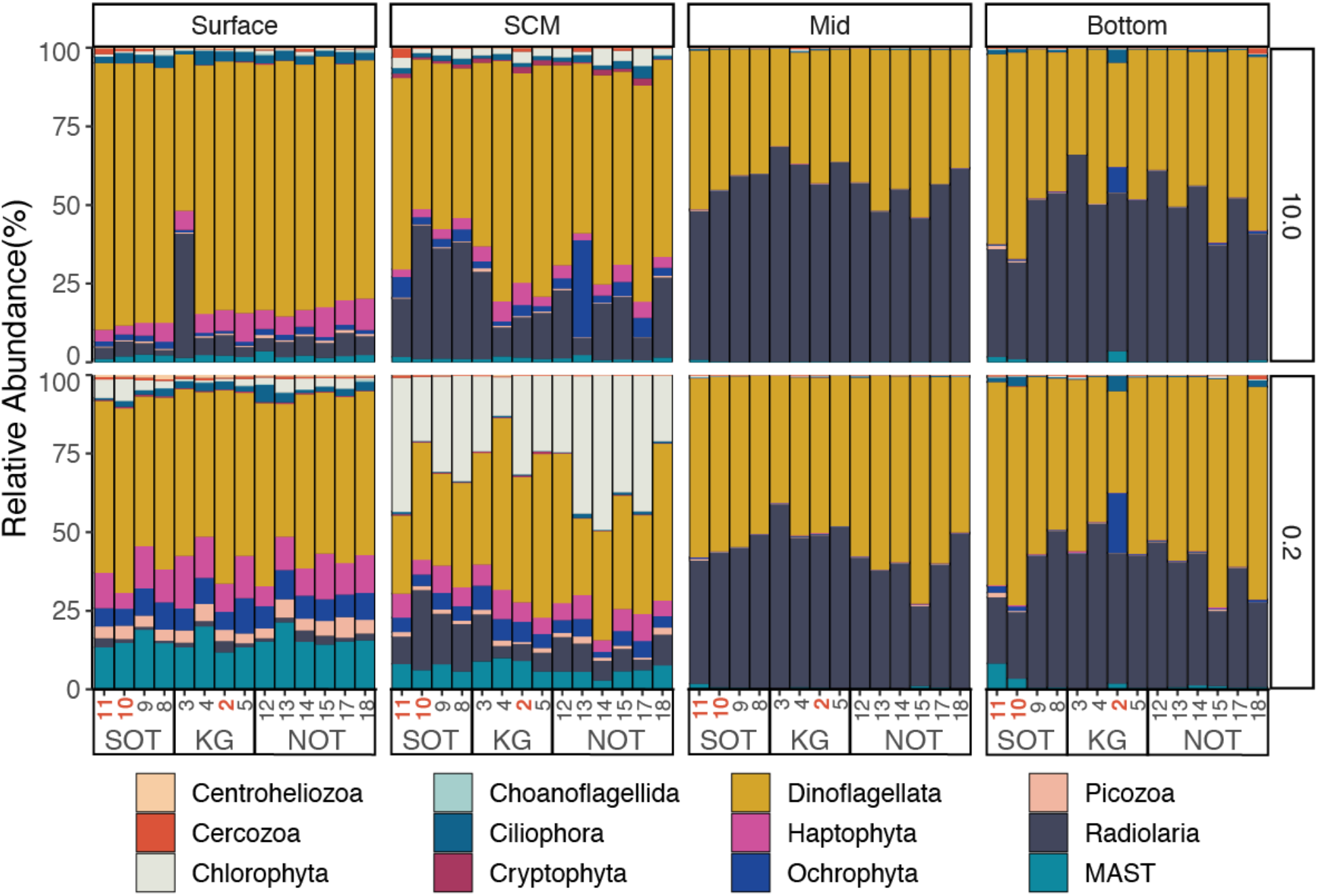
Relative abundance of major protist groups in size-fractionated samples from four depths in the Okinawa Trough (OT). Sampling stations are ordered on the x-axis from south to north: Southern Okinawa Trough (SOT) stations include 11, 10, 9, and 8; Kerama Gap (KG) stations include 3, 4, 2, and 5; Northern Okinawa Trough (NOT) stations include 12, 13, 14, 15, 17, and 18. The plot is faceted by sampling depth (columns) and filter pore-size in μm (rows). When replicates were available for a particular station/depth/filter combination, replicates were collapsed and represented with a single stacked bar. Regional community differences are not visible at the high taxonomic level represented in the plot, but there are visible differences by depth and filter pore-size. Hydrothermally influenced sites include station 10 at the Hatoma Knoll vent field, station 11 near the Dai-Yon Yonaguni Knoll, and station 2 at the ANA vent at the Daisan-Kume Knoll (highlighted in red).

### Protist biogeography in the Okinawa Trough

To investigate biogeography of protist communities in the OT, we merged smaller and larger size-fractions for replicates and subset samples by depth. Permutational Multivariate Analyses of Variance (PERMANOVA, 999 permutations) by region (SOT, KG, NOT) on the Aitchison distances between samples were significant in surface, SCM, mid-water, and bottom-water samples (*p* < 0.05). Among surface samples, NOT and SOT samples were significantly different from KG samples, but NOT and SOT samples were not significantly different from each other. Among SCM samples, NOT and SOT samples were significantly different, but neither were significantly different from KG samples. For mid- and bottom-water samples, all pairwise comparisons were significant.

Redundancy analysis was applied to evaluate the contribution of environmental factors to regional differences in protist community composition (Figure 4). ANOVA results for the surface water RDA model were significant (*p* < 0.05) and ANOVA by environmental variable showed that temperature significantly affected community composition. Variance partitioning indicated that temperature explained 2.6% of variance, whereas region explained 5.5% (multiple testing adjusted R^2^). Dissolved Oxygen (DO) was correlated with temperature (Supplemental Figure 3) in surface samples, so DO was excluded from the model. The RDA model for SCM samples additionally included the depth of the SCM at each site and no variables covaried. The SCM model results were not significant (*p* = 0.182); variance partitioning indicated region explained 3.4% of variance. In the mid-water samples, temperature covaried with DO, CDOM, nitrate, silicate, and phosphate and, of these, only temperature was included in the RDA model. The mid water model results were statistically significantly and salinity and NH_4_ both significantly contributed to model results. Variance partitioning showed that region explained 7.3% of variance, whereas NH_4_ and salinity explained 2.4 and 1.2%, respectively. In bottom water samples, temperature covaried with CDOM, nitrate, phosphate, silicate and, of these, only temperature was included in the RDA model. The RDA model was statistically significant and turbidity, NH_4_, temperature and DO significantly contributed to model results. Variance partitioning indicated that region explained 6.9% of variance, whereas turbidity explained 3.2%, NH_4_ explained 1.4%, temperature explained 1.5% and DO explained 1.7 %. It should be noted that DO and temperature had very high variance inflation factors in this test (> 150), even though their correlation coefficient was < 0.8.

**Figure 4.**
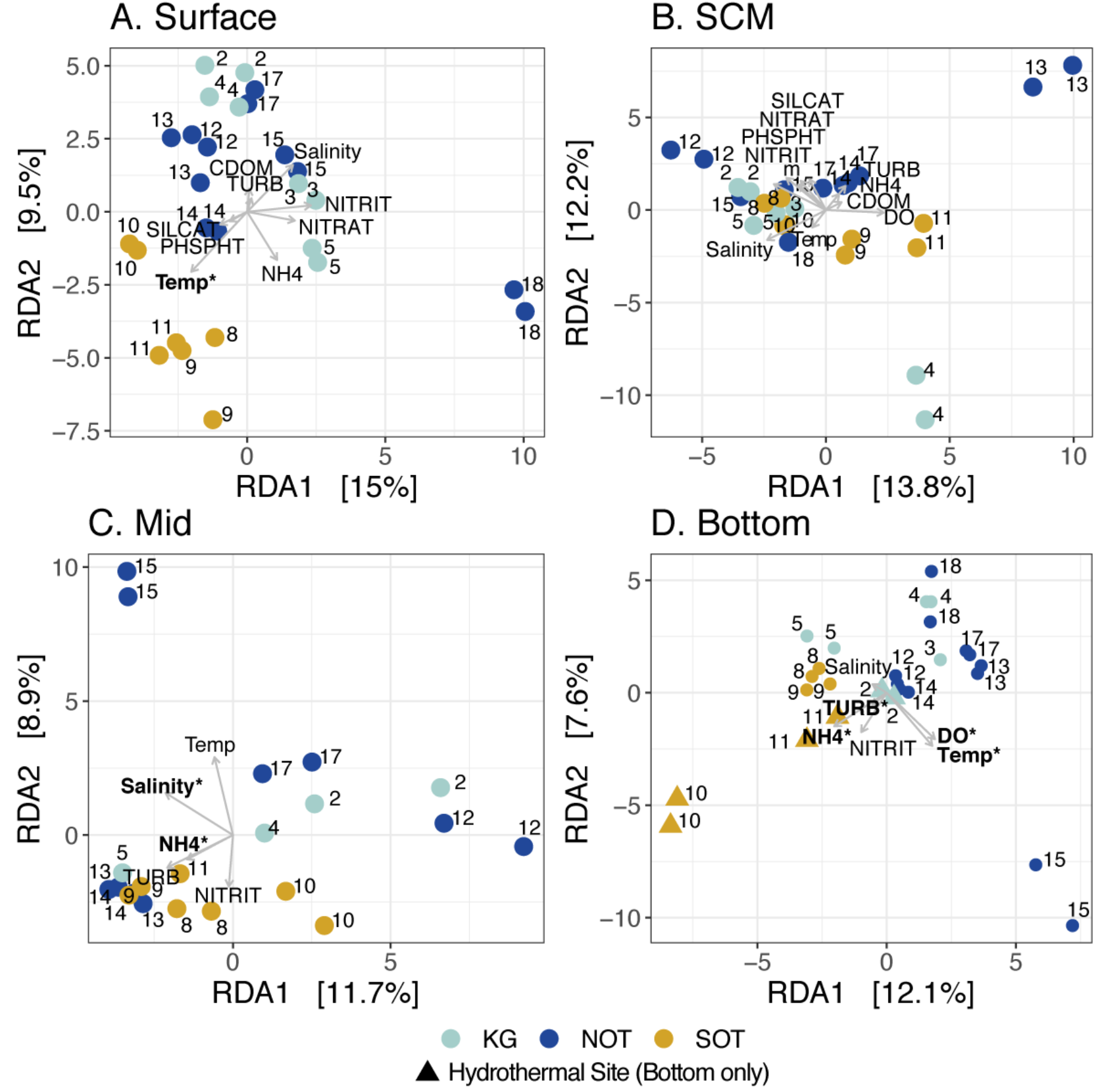
Distance based redundancy analysis (RDA) of protist communities and explanatory environmental variables for four depth-layers in the Okinawa Trough. If environmental variables covaried with a Pearson correlation coefficient greater than 0.8 (absolute value), only one covarying variable was included in the analysis; temperature covaried with dissolved oxygen (DO) in surface waters, no variables covaried at the SCM, temperature covaried with DO, colored dissolved organic matter (CDOM), nitrate (NITRAT), silicate (SILICAT), and phosphate (PHSPHT) in the mid-water column, and temperature covaried with CDOM, nitrate, phosphate, silicate in the bottom waters. The RDA model results were significant in surface (A), mid (C), and bottom waters (D) and variables that significantly contributed to variation in community composition are in bold and marked with an asterisk (ANOVA, *p* < 0.05). KG is the Kerama Gap (light blue), NOT is the Northern Okinawa Trough (dark blue), and SOT is the Southern Okinawa Trough (gold). For bottom water samples (D), hydrothermally influenced sites are demarcated as triangles.

### Protist communities at hydrothermal vent sites

To evaluate the effect of hydrothermal activity on protist diversity, we first checked whether alpha diversity was significantly different at hydrothermally influenced sites; hydrothermal influence did not significantly increase or decrease observed richness or Shannon indices in bottom water samples (Supplemental Figure 4C–D). To determine if hydrothermal influence altered community composition, we performed PERMANOVA by presence or absence of hydrothermal influence on the Aitchison distances between bottom water samples; community compositions at hydrothermally influenced sites were significantly different from other bottom water samples (*p* = 0.001).

Lastly, we investigated if there were protist ASVs specifically associated with hydrothermally influenced sites by testing for differentially abundant ASVs in the hydrothermally influenced bottom samples compared to the rest of the bottom water samples using DESeq2 [37]. An ASV was considered significantly differentially abundant when the False Discovery Rate adjusted *p*-value was less than or equal to 0.01. Overall, 45 ASVs were significantly differentially abundant between samples from hydrothermally influenced sites and the rest of the bottom water samples. Of the differentially abundant ASVs, 30 were more abundant at sites with hydrothermal influence than at the other sites, and included Dinophyceae, Syndiniales, Chrysophyceae, MAST, RAD-A and RAD-B Radiolarians, Oligohymenophorea and Spirotrichea ciliates, and Picozoa ASVs (Figure 5).

**Figure 5.**
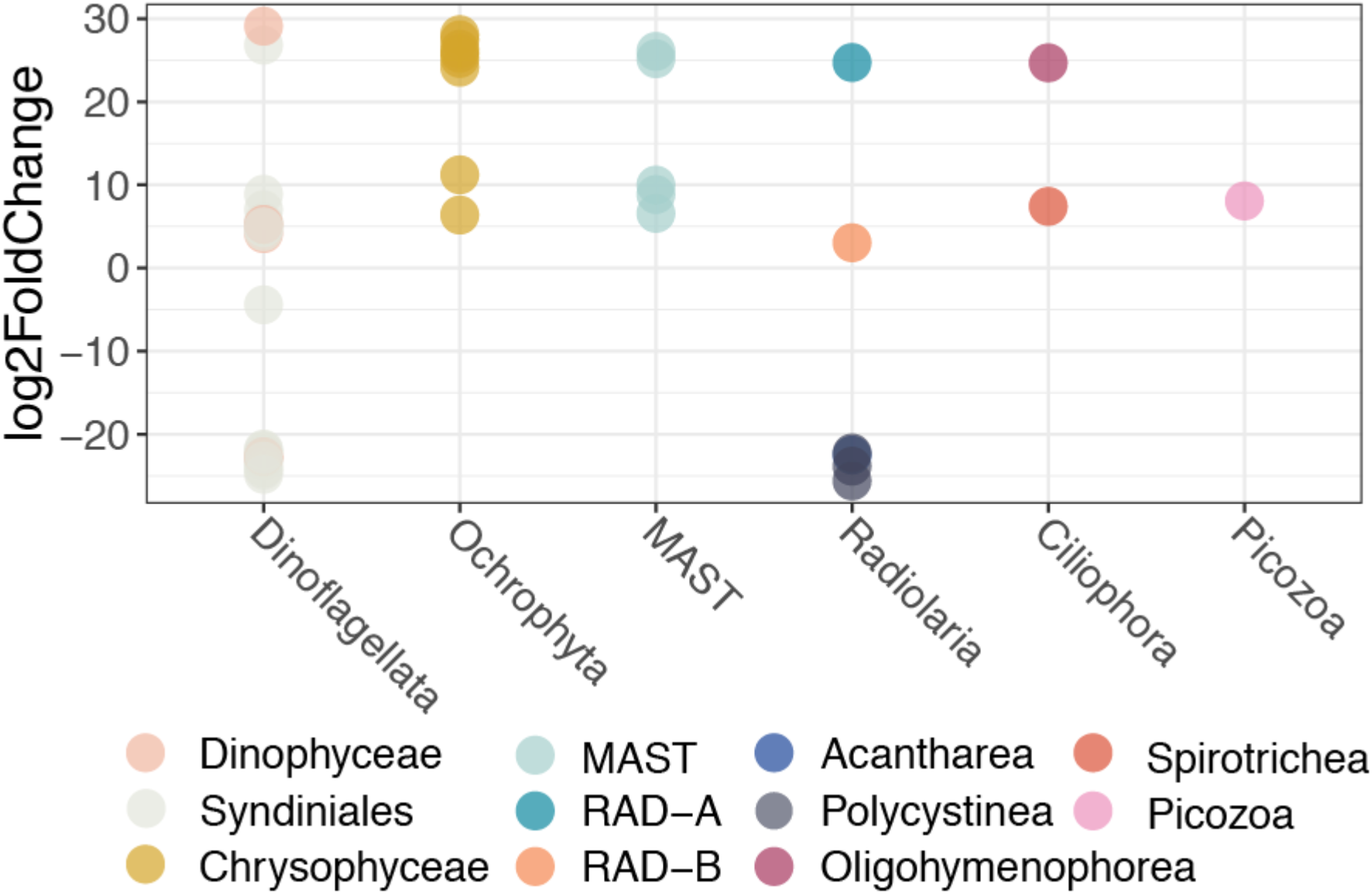
Log2 fold-change of significantly differentially abundant ASVs in bottom water influenced by hydrothermal activity. The DESeq function in the DESeq2 package was used to test whether ASVs were significantly more (positive log2 fold-change) or less (negative log2 fold-change) abundant in hydrothermally influenced bottom water samples from Stations 10, 11, and 2 compared to bottom water samples from the other sites. Each point represents a single significantly differentially abundant ASV and taxonomic groupings are indicated on the x-axis and by color. ASVs were considered significantly differentially abundant if False Discovery Rate adjusted *p*-values were < 0.01.

## Discussion

Due to their small size, protists, like bacteria, face few dispersal barriers in the ocean [38]. In addition, rapid asexual reproduction in many protists, as well as the ability to form resting cysts, increases their dispersal potential. As a result, many protists are cosmopolitan, suggesting that global protist diversity is low while local diversity in any given place is relatively high [39, 40]. Whether or not this is the case depends largely on how protist species and diversity are defined. For example, if a higher level of genetic diversity is included in a species definition, global diversity will be lower. Conversely, if less genetic differentiation is included in a species definition, global diversity will be higher [41]. This has been demonstrated in bacteria through studies of biogeography at deep sea hydrothermal vents: only when microdiversity—diversity at the single nucleotide polymorphism level—is considered does geographic population structure emerge [17, 42]. By analyzing ASVs, which capture the highest amount of diversity available in metabarcoding datasets, we investigated whether biogeographic patterns exist among protists in the Okinawa Trough in regard to regional ocean circulation patterns, water mass composition, and hydrothermal influence. Our results demonstrate relatively weak biogeography in surface waters through to the mid-water column, but show strong regional differentiation of protist communities in deeper water. In addition, protist communities in bottom waters affected by hydrothermal vent activity were significantly different from communities in other bottom water samples.

### Protist biogeography in the Okinawa Trough

Consistent with previous studies on protist biogeography, the strongest overall influence on community composition was the depth sampled [43]. Depth-dependent diversity patterns were similar to metabarcoding results from other regions, with higher diversity in the surface and SCM than the meso- and bathypelagic [8, 43]. In addition, overall community structure by depth was comparable to global patterns: Dinoflagellata, Haptophyta, Ochrophyta, Picozoa, Radiolaria and Marine Stramenopiles (MASTs) were abundant in the surface and SCM while Dinoflagellata and Radiolaria dominated in the meso- and bathypelagic [5].

While depth was the strongest determinant of community composition overall, there were significant regional differences in community composition in all depth layers and region contributed more to variance than environmental variables. The Okinawa Trough is divided into Southern (SOT) and Northern (NOT) regions based on the location of the Kerama Gap (KG) and bottom water characteristics [10]. The fast-flowing, northward Kuroshio current is the major oceanographic feature in the upper water column [9]. Intermediate water consistently enters the trough through the KG, whereas surface water regularly enters and occasionally exits [11, 44]. Regional differences in protist community composition are mostly consistent with these prevailing current patterns. In surface and mid-water samples, NOT and SOT communities were both significantly different from KG samples, but NOT and SOT communities were not different from each other in the surface waters, where the Kuroshio current is strongest. At the SCM, NOT and SOT samples differed significantly but KG samples did not differ from NOT or SOT samples. These results may be due, in part, to Changjiang (a.k.a. Yangtze) Diluted Water (CDW) and Kuroshio water were mixing at the continental shelf near station 13. The Changjiang/Yangtze is the longest river in China and transports terrestrial nutrients and anthropogenic pollutants into the East China Sea, causing eutrophication that regularly triggers causes diatom blooms [45]. CTD and nutrient measurements at station 13 showed decreased subsurface salinity and increased nutrient concentrations and chlorophyll fluorescence compared to other sampling sites (Supplemental Figure 6). In addition, there as was an increased contribution of Bacillariophyceae diatoms (Ochrophyta, Figure 4) to the community composition at station 13.

The regional pattern in protist community was most robust in the bottom waters, where protist communities were significantly distinct in each region and where each region has distinct water mass composition; bottom water in the SOT is comprised of 45% low-salinity North Pacific Intermediate Water (NPIW) and 55% high-salinity South China Sea Intermediate Water (SCSIW), while KG bottom water is 75% NPIW and 25% SCSIW and NOT bottom water is 65% NPIW, 35% SCSIW [10]. Similarly, Agogue et al. (2011) found water mass composition was the strongest predictor of prokaryotic community composition in the Atlantic Ocean [6], and Pernice et al. (2016) found water mass composition was the strongest predictor of protist community composition in the deep waters of the Atlantic, South and Equatorial Pacific, and Indian oceans [5]. Water masses in the ocean may either act as geographic boundaries to dispersal or select for microbes adapted to specific environmental conditions [5, 6]. If differences in community composition were due to distinct chemical properties of water masses, it would support the latter [6]. In the OT, salinity is the major marker for water mass composition [10], and the salinities measured during our sampling were consistent with those reported for the different water mass compositions (Supplemental Figure 2A–B, [10]). However, salinity did not significantly contribute to variation in community composition in the bottom waters of the OT (Figure 4D). Therefore, our results support the idea that particular water mass histories influence community composition and suggest that water masses may act as barriers to microbial dispersal [6].

### Protist communities at hydrothermal vent sites

The protist communities in samples from hydrothermally influenced sites were significantly different from the communities in bottom waters at sites without hydrothermal influence. Furthermore, environmental factors that significantly contributed to variation in community composition in bottom waters—turbidity, NH_4_—are related to hydrothermal activity in the OT, where vents resuspend overlying terrigenous material creating turbid vent plumes [16]. Syndiniales, Ciliophora, and MAST (Marine Stramenopiles) ASVs were significantly enriched in hydrothermally influenced OT bottom waters, which is consistent with previous work comparing protists in bottom waters with varying proximity to hydrothermal vents [3]; Ochrophyta, environmental clades of Radiolaria, and Picozoa ASVs were additionally enriched near hydrothermally influenced sites in this study (Figures 2, 5). Quantitative models of deep-water circulation suggest that vents in back-arc basins, such as the Okinawa Trough, are well-connected, but basin to basin dispersal may be restricted [12]. A remaining question, then, is whether protists enriched near vents in the Okinawa Trough are similar to protists in other hydrothermal systems and other marine ecosystems.

Dinoflagellate ASVs made up the majority of differentially abundant ASVs at hydrothermally influenced sites (22 out of 45, Figure 5) and half of these were more abundant near vents. Among the enriched ASVs, eight are in the parasitic order Syndiniales, including group I (clades 2 and 7) and group II (clades 1, 6, 12, 13, 14, and 16). The remaining enriched dinoflagellate ASVs belong to the Order Dinophyceae but could not be further classified. Group II Syndiniales are common in sunlit, surface waters, whereas group I is common in suboxic and anoxic ecosystems. However, both groups I and II are found in surface and deep water and have clades that have only been recovered from suboxic or anoxic ecosystems [46]. The majority of the enriched Syndiniales ASVs shared 100% identity with GenBank sequences (nr/nt, accessed 5/16/2019, [47, 48]) from a wide variety of ecosystems throughout the global ocean, including surface waters near the poles, mesopelagic water near the California coast, 100 and 200 m oxygenated and micro-oxic North Atlantic waters, and at the sediment interface near the Juan de Fuca Ridge vent system [49, 50]; the Syndinales ASVs enriched in hydrothermally influenced samples clearly derive from cosmopolitan organisms. Although they are not restricted to vent systems, parasitic Syndiniales do seem to thrive in vent systems and may take advantage of increased host availability in such regions [51].

Ochrophyta accounted for the second-most enriched ASVs in hydrothermally influenced samples (Figure 5). The enriched Ochrophyta ASVs belong to several clades within the order Chrysophyceae. Two belong to clades with unknown morphology made up of environmental sequences (clades EC2H and EC2I) recovered from a diverse array of marine and freshwater environments [52]. The remaining ASVs belong to the genus *Paraphysomonas*, which are colorless phagotrophs that are globally distributed in freshwater, marine, and soil ecosystems and have previously been recovered from vent sites [53]. Although several of the enriched ASVs share > 99%–100% identity with previously published vent-associated sequences [53], they are equally similar to sequences recovered from many other marine environments, indicating that these enriched ASVs also derive from cosmopolitan organisms.

Five of the ASVs enriched in hydrothermally influenced samples belong to marine stramenopile lineages—MAST-1, −7, −8 (Figure 5). Most of what is known about MAST lineages has been learned through environmental molecular surveys. Aggregate analyses of sequences collected through such studies have found that MAST-1, −7, and −8 are highly diverse, globally-distributed, and abundant in surface waters [54, 55]. Culture-independent techniques have further demonstrated that MAST-1 [56], MAST-7 [57], and MAST-8 [54] are bactivorous flagellates. Therefore, it is likely that the MAST ASVs enriched in hydrothermally influenced samples represent cosmopolitan bacterivores. Indeed, MAST ASVs from this study each share 99% (MAST-7) to 100% (MAST-1, −8) identity with multiple sequences recovered from surface waters, including from (anoxic) Saanich Inlet, Vancouver, Canada [58], the Scotian Shelf in the North Atlantic [59], and the Southern Ocean [60].

The RAD-A and RAD-B radiolarian groups represent environmental clades that have no morphological description—other than being picoplankton—or known ecological roles [61]. The RAD-A ASV enriched in hydrothermally influenced samples (Figure 5) shares > 99% identity with many GenBank sequences recovered from oxic and anoxic waters globally, including the Cariaco basin in the Caribbean Sea [50], the Gulf Stream [49], and the South East Pacific [62]. The enriched RAD-B ASV shares 100% identity with multiple sequences recovered from the East Pacific Rise (1500 m and 2500 m), the Arctic Ocean (500 m), oxygenated water in the Cariaco Basin [50], the Juan de Fuca Ridge [63], and the Southern Ocean [64]. While the radiolarian ASVs enriched in hydrothermally influenced samples appear to be cosmopolitan, it is notable that identical and similar sequences have repeatedly been isolated from vent fields and anoxic regions.

Previous studies indicated Ciliophora are abundant in sediments and bacterial mats near hydrothermal vents [65–67]. It is unsurprising, then, that two Ciliophora ASVs were significantly more abundant in hydrothermally influenced samples (Figure 5). The enriched Spirotrichea ASV was classified to genus level by our classifier and belongs to Leegaardiella, a recently described genus collected in the North Atlantic [68]. The Leegardiella ASV shared 100% coverage and identity with one sequence in GenBank— an uncultured eukaryote clone recovered from 2500 m at the East Pacific Rise—and had greater than 99% shared identity with another sequence recovered from the East Pacific Rise, also from 2500 m [49]. However, the Leegardiella ASV also had 100% coverage and > 99% identity matches with several sequences recovered from Arctic surface water [69]. This Leegardiella ASV may, therefore, derive from a widely distributed genus that opportunistically becomes more abundant at vent sites. The Oligohymenophorea ASV belongs to the environmental OLIGO 5 clade, but did not have > 96% identity matches to any sequences in GenBank; the closest match was to an uncultured eukaryote clone recovered from 500 m depth off the coast of California [49]. The EukRef-Ciliophora curated Oligohymenophorea phylogeny groups the OLIGO 5 clade with the subclass Scuticiliatia [70]. Interestingly, Zhao and Zu (2016) also found Spirotrichea and Scuticiliatia enriched among ciliates found near a vent site in the Northern Okinawa Trough [71]. While spirotrich ciliates are generally bacterivores [65], Scuticiliatia ciliates are known to be symbionts of aquatic organisms [72], including giant hydrothermal bivalves [3]. If the Scuticiliata ASV is a symbiont, it could provide some explanation as to why more similar sequences have not been recovered elsewhere; coevolution with a dispersal-limited host could also limit symbiont dispersal [73]. However, it is equally likely that sequences with high similarity to the Scuticiliata ASV have not yet been recovered elsewhere simply due to insufficient sampling.

A single ASV enriched near vent sites belonged to Picozoa (Figure 5). The first Picozoa species was described in 2013 [74] and Picozoa ecology remains poorly resolved. Information regarding their distribution derives from environmental molecular surveys, which have established five clades within Picozoa. While all Picozoa are marine, biogeographical patterns vary between clades: clade BP2 has only been found in surface waters, group 2 is mostly found in deep waters, and the other three groups are globally distributed throughout the water column [75]. The enriched Picozoa ASV belongs to one of the cosmopolitan clades (group 1). The sequence shares 100% identity with three others in Genbank, from a variety of environments—surface water in the Arctic [76] and Southern [77] oceans, as well as mesopelagic water near southern California [49].

### Conclusions and future directions

While the Okinawa Trough provides an ideal setting to investigate protist diversity and biogeography, this study represents the first comprehensive survey of protists in the region. Our results show that protist communities largely reflect regional oceanography and are influenced by water mass composition in near-bottom waters. Disentangling environmental and geographic effects on communities is challenging since many variables, such as temperature, covary with latitude. However, region consistently contributed more to variance in community composition than any environmental factors in this study. We additionally found distinct protist communities associated with hydrothermally influenced sites, with bacterivore, parasitic, and putatively symbiotic protists enriched in hydrothermally influenced bottom waters. The enriched protists were mostly globally distributed, and many were previously found at vent sites in other parts of the world. These results emphasize the importance of the rare biosphere; most protists seem to be widely distributed, even if rare, and can become opportunistically more abundant under certain conditions. A major limitation of metabarcoding studies with DNA is that it is impossible to know whether sequences derive from actively metabolizing cells, dead or inactive cells, or even environmental DNA; future studies will benefit from incorporating methods that better delineate metabolic state. Additionally, metabarcoding studies suffer from focusing on a short sequence to assess diversity; metagenomic and metatranscriptomic approaches are becoming increasingly feasible for large-scale protist surveys, which will allow for finer resolution of diversity. In the future, comparisons between the Okinawa Trough and other hydrothermal regions will continue to advance our understanding of important protists in hydrothermal ecosystems and how hydrothermal activity influences protist diversity and biogeography in the global ocean.

## Acknowledgements

We thank the captain and crew of the JAMSTEC R/V Mirai for their assistance and support in sample collection. Hiroyuki Yamamoto, Hiromi Watanabe, Dhugal Lindsay, Mary M. Grossmann, and Yuko Hasagawa were instrumental in organizing and facilitating cruise sampling and Otis Brunner substantially contributed to seawater filtration. We are further grateful to Hiromi Watanabe for helping to access and interpret environmental data and for comments on the manuscript. We thank the OIST DNA sequencing section (Onna, Okinawa) for carrying out the sequencing. This work was funded by the Marine Biophysics Unit of the Okinawa Institute of Science and Technology Graduate University. MMB was supported by a Japan Society for the Promotion of Science DC1 graduate student fellowship.

## Notes

#### Summary of Updates

Additional analyses including environmental data have been added.

https://github.com/maggimars/OkinawaTroughProtists

https://maggimars.github.io/OkinawaTroughProtists/OTprotists.html

