## Supplemental Figures for "Regional oceanographic features and hydrothermal activity influence protist diversity and biogeography in the Okinawa Trough"

**Supplemental Table:**
**Supplemental Table 1.** Coordinates, sampling depths, and total depth for all sampling stations.

**Supplemental Figures:**
**Supplemental Figure 1.** Depth profiles of colored dissolved organic matter (CDOM) (µg/L), turbidity (FTU), NH4 (µmol/kg), and total carbon (µmol/kg)
**Supplemental Figure 2.** Cross-section of salinity in the Kerama Gap (A) and bottom-water salinity values throughout the Okinawa Trough and Kerama Gap (B).
**Supplemental Figure 3.** Heatmap of Pearson correlation coefficients (absolute values) between environmental and regional variables at four depths in the Okinawa Trough.
**Supplemental Figure 4.** Alpha diversity (observed richness and Shannon indices) for protist communities in size-fractionated samples from the Okinawa Trough.
**Supplemental Figure 5.** Principal coordinates analysis of Aitchison distances between size-fractionated protist communities from four depth layers in the Okinawa Trough.
**Supplemental Figure 6.** Depth Profiles for fluorescence (µg/l), salinity (PSU) and nutrient concentrations (µmol/kg) in the top

**Supplemental Table 1. Coordinates, sampling depths, and total depth for all sampling stations.**

| Station | Longitude (°E) | Latitude (°N) | SCM Depth (m) | Mid Depth (m) | Bottom Depth (m) | Site Depth (m) |
|---|---|---|---|---|---|---|
| 2 | 126.468 | 26.290 | 92 | 700 | 1066 | 1080 |
| 3 | 127.500 | 25.416 | 72 | 1000 | 2385 | 2407 |
| 4 | 126.900 | 25.928 | 58 | 700 | 1834 | 1857 |
| 5 | 126.084 | 26.501 | 82 | 700 | 1900 | 1922 |
| 8 | 124.994 | 25.942 | 88 | 700 | 1671 | 1681 |
| 9 | 124.012 | 25.502 | 74 | 700 | 1890 | 1914 |
| 10 | 123.837 | 24.857 | 85 | 700 | 1515 | 1530 |
| 11 | 122.840 | 24.760 | 50 | 700 | 1217 | 1223 |
| 12 | 126.903 | 27.785 | 80 | 700 | 1024 | 1033 |
| 13 | 127.339 | 29.003 | 80 | 700 | 834 | 846 |
| 14 | 127.501 | 28.747 | 75 | 700 | 1013 | 1025 |
| 15 | 129.029 | 28.792 | 67 | 700 | 776 | 783 |
| 17 | 129.570 | 28.957 | 90 | 700 | 772 | 779 |
| 18 | 130.904 | 28.981 | 100 | 1500 | 2957 | 2981 |

**A.**

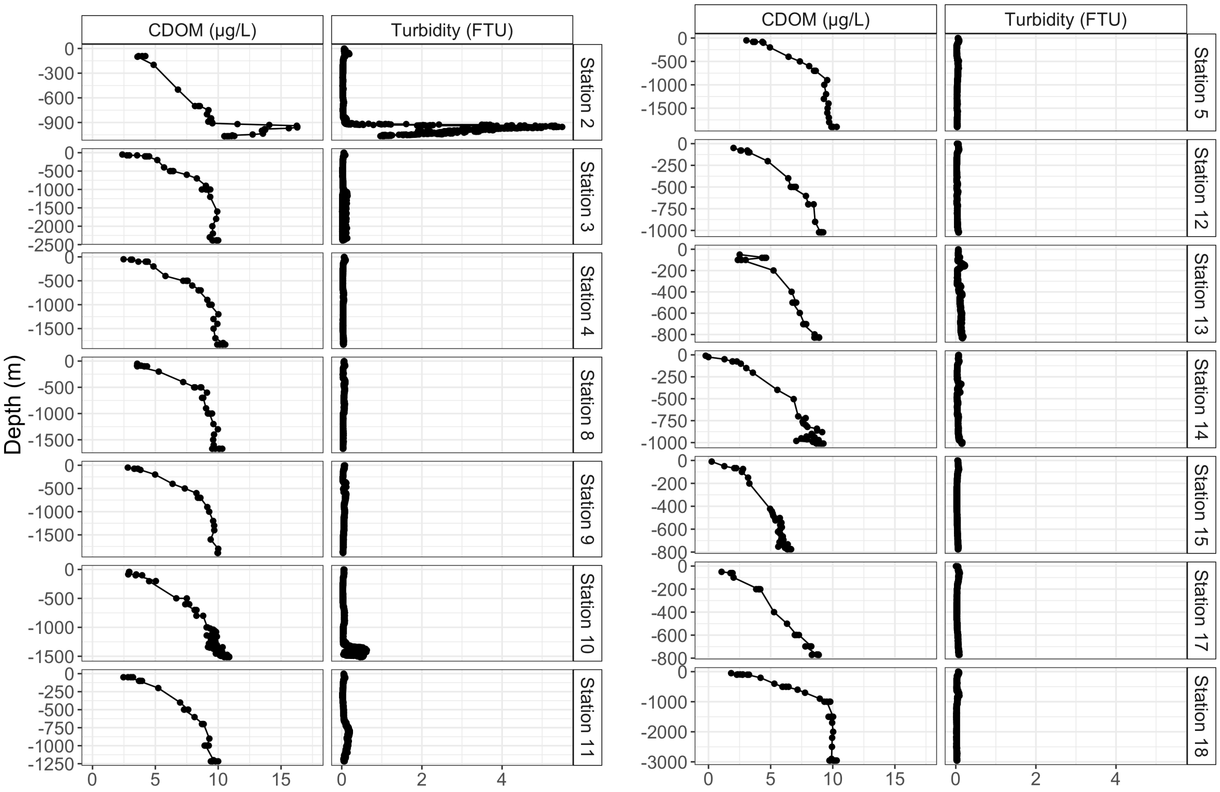

CDOM (μg/L)
Turbidity (FTU)
Depth (m)
Station 2
Station 3
Station 4
Station 8
Station 9
Station 10
Station 11
Station 5
Station 12
Station 13
Station 14
Station 15
Station 17
Station 18

**B.**

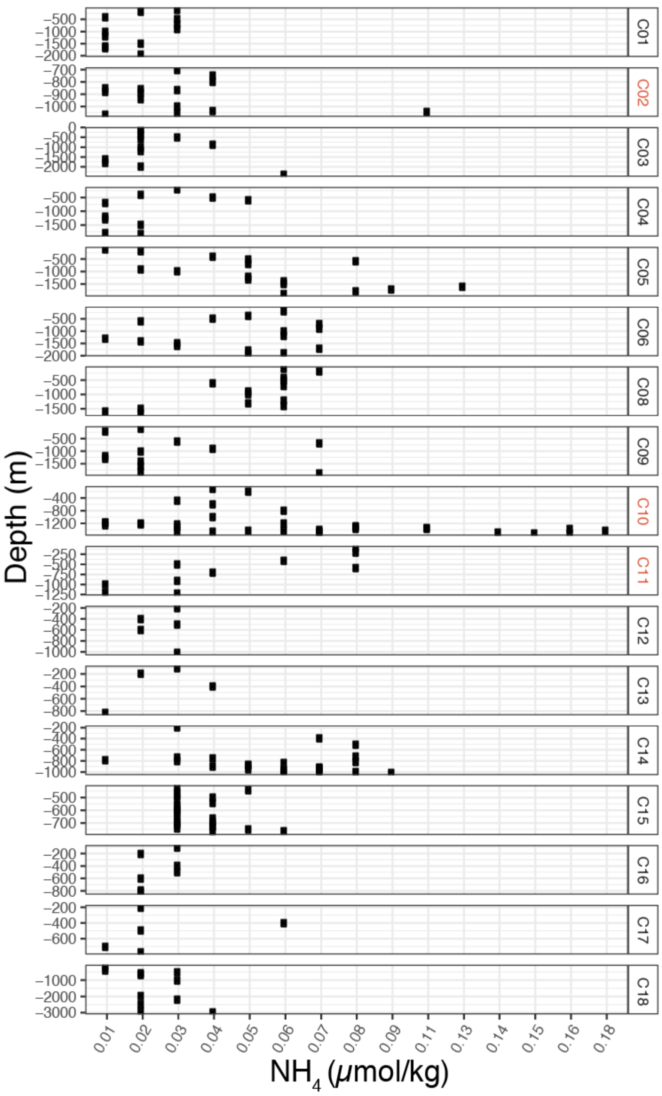


**C.**

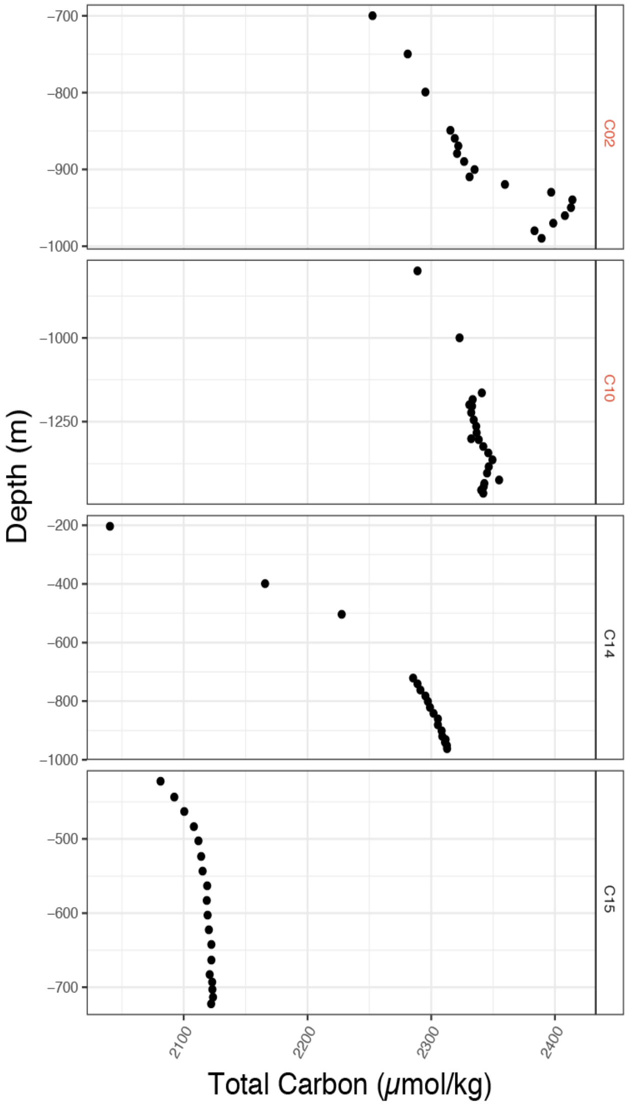


**Supplemental Figure 1. Depth profiles for colored dissolved organic matter (CDOM) (µg/L), turbidity (FTU), NH4 (µmol/kg), and total carbon (µmol/kg).** CDOM measurements were made on water collected in Niskin bottles during the Niskin Rosette and CTD cast at each station; turbidity was measured directly by CTD sensor during the same casts. Deep CDOM and turbidity maxima, which indicate hydrothermal activity in the Okinawa Trough [20], are visible in profiles from stations 2, 10, and 11, but not at other stations.

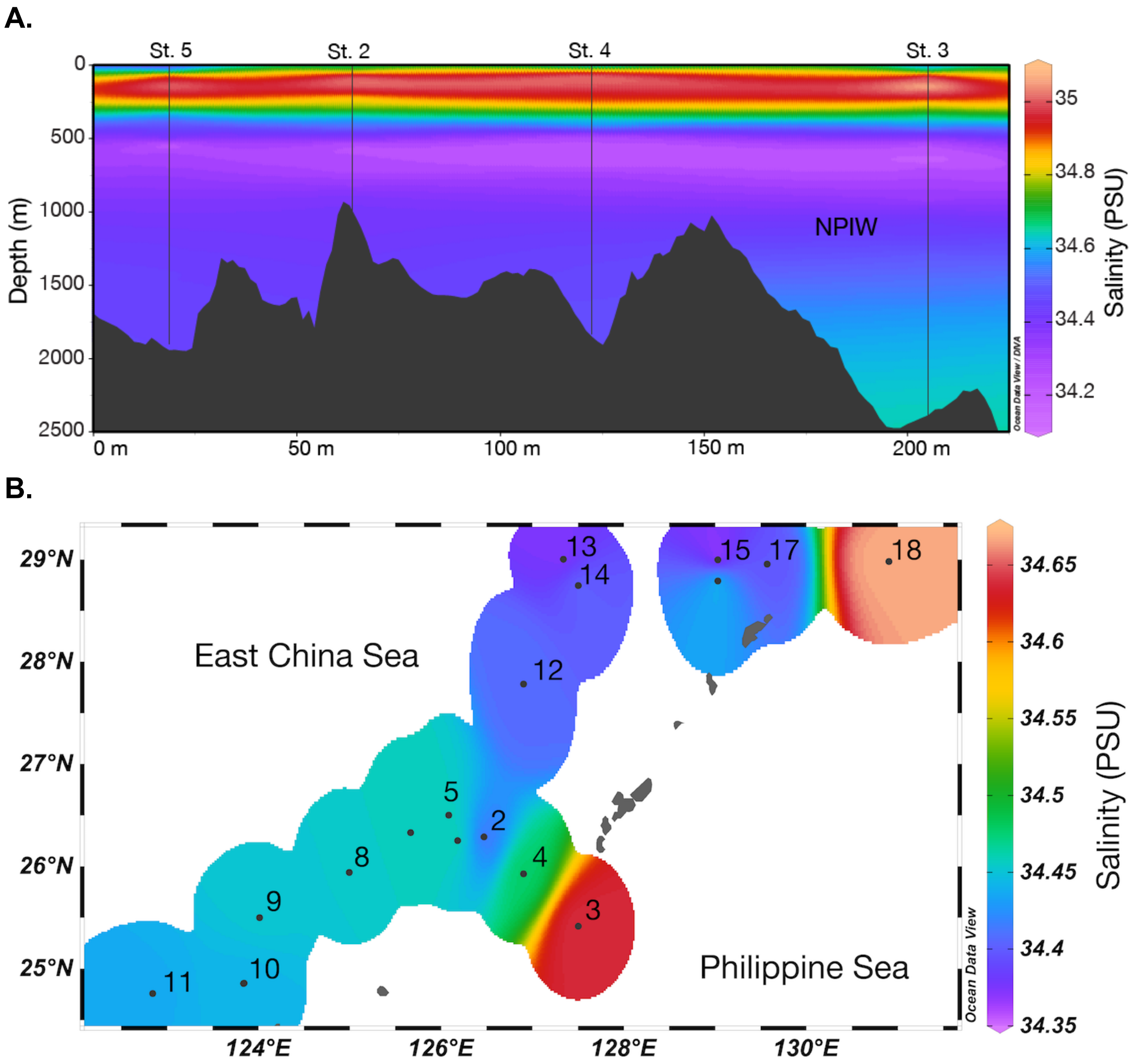


**Supplementary Figure 2. Cross-section of salinity in the Kerama Gap (A) and bottom-water salinity values throughout the Okinawa Trough and Kerama Gap (B).** Salinities were plotted and extrapolated from CTD measurements with Ocean Data View weighted-average gridding (http://odv.awi.de). Low-salinity North Pacific Intermediate Water (NPIW) flows through the Kerama Gap into the Okinawa Trough and contributes to the lower salinity in bottom waters of the Northern Okinawa Trough compared to the southern trough.

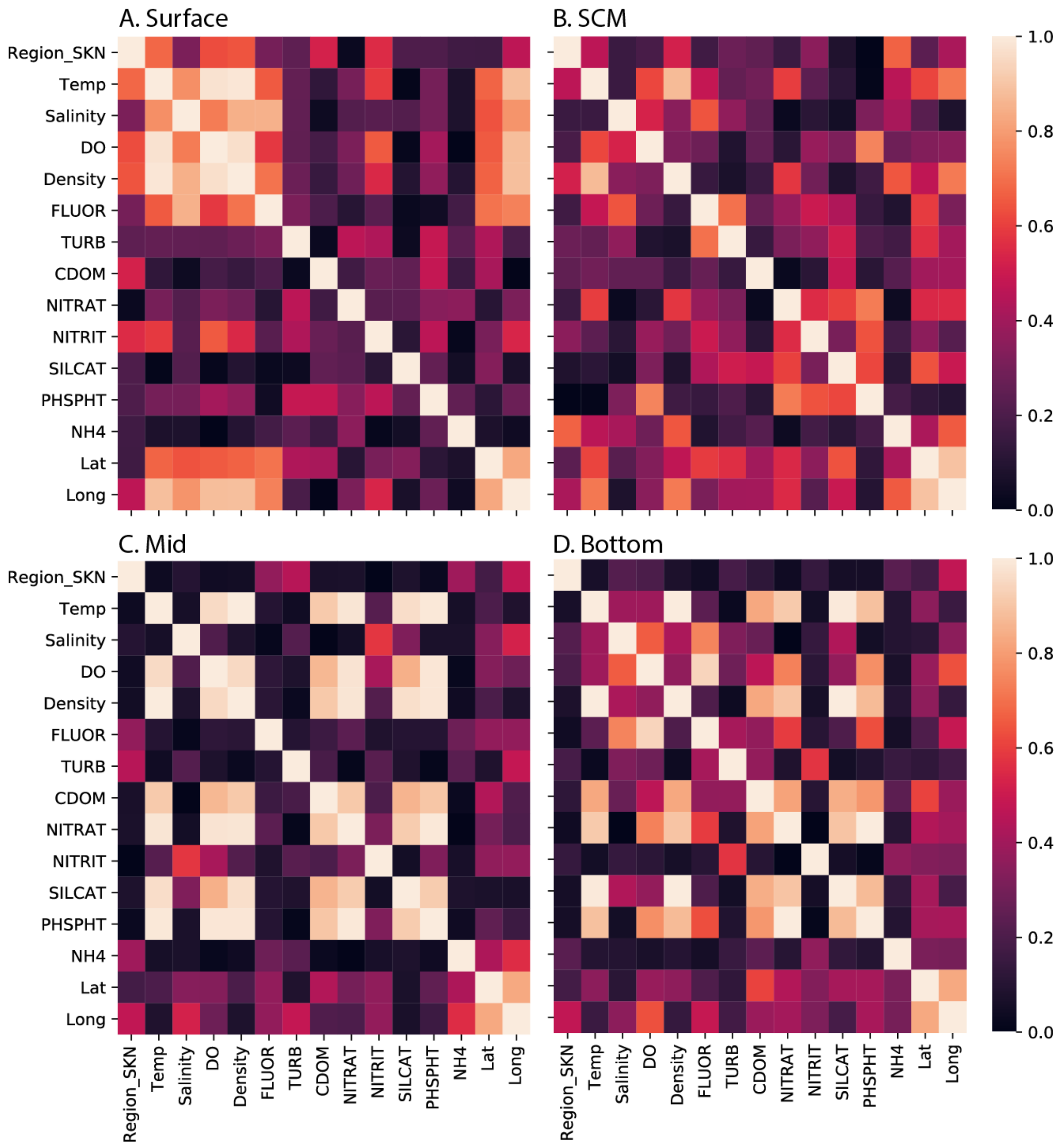


**Supplemental Figure 3. Heatmap of Pearson correlation coefficients (absolute values) between environmental and regional variables at four depths in the Okinawa Trough.** Correlation coefficients were computed and plotted with the python packages pandas and seaborn. If absolute values of correlation coefficients exceeded 0.8, only one of the correlated variables was included in redundancy analysis and variance partitioning for that depth.

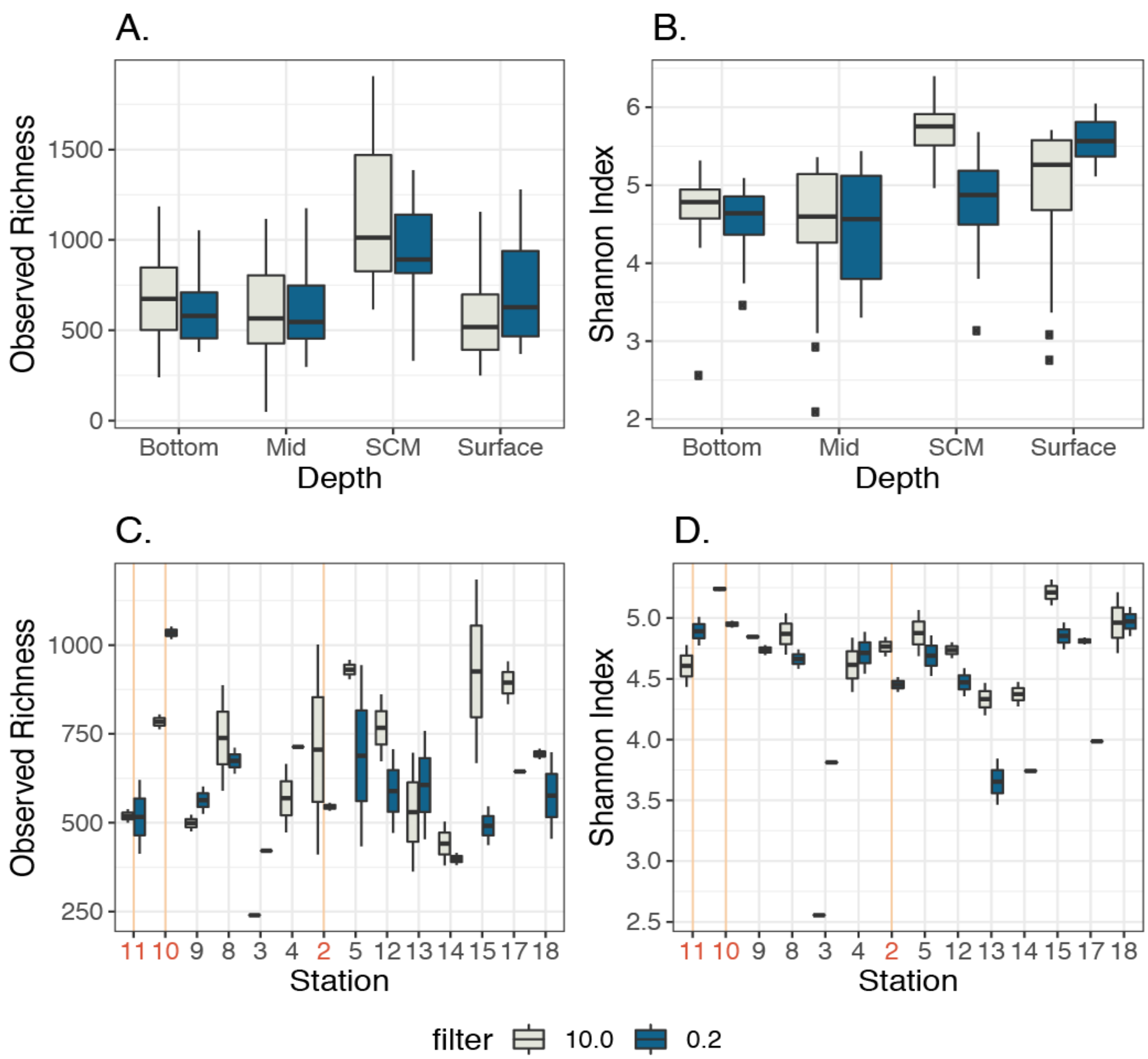


**Supplemental Figure 4. Alpha diversity for protist communities in size-fractionated samples from the Okinawa Trough**. A. Observed Richness and B. Shannon Indices in samples collected from four depth layers at 14 stations in the Okinawa Trough. C. Observed Richness and D. Shannon Indices in samples collected from near-bottom water. In C and D, stations 11, 10, and 2 are highlighted to indicate sites with hydrothermally influenced bottom waters. Boxes depict upper and lower quartiles; whiskers depict ranges and points are outliers. Tan-boxes represent the larger size-fraction (> 10 µm) and blue boxes represent smaller size-fractions (< 10 µm, > 0.2 µm). SCM samples have significantly higher richness than surface, mid, and bottom water samples ($p < 0.001$). Shannon indices for SCM and surface samples were not significantly different but were both significantly higher than for mid and bottom water samples. ($p < 0.001$). Hydrothermally influenced bottom waters at stations 11, 10, and 2 (highlighted in panels C and D) did not have significantly different observed richness or Shannon indices from other bottom water samples.

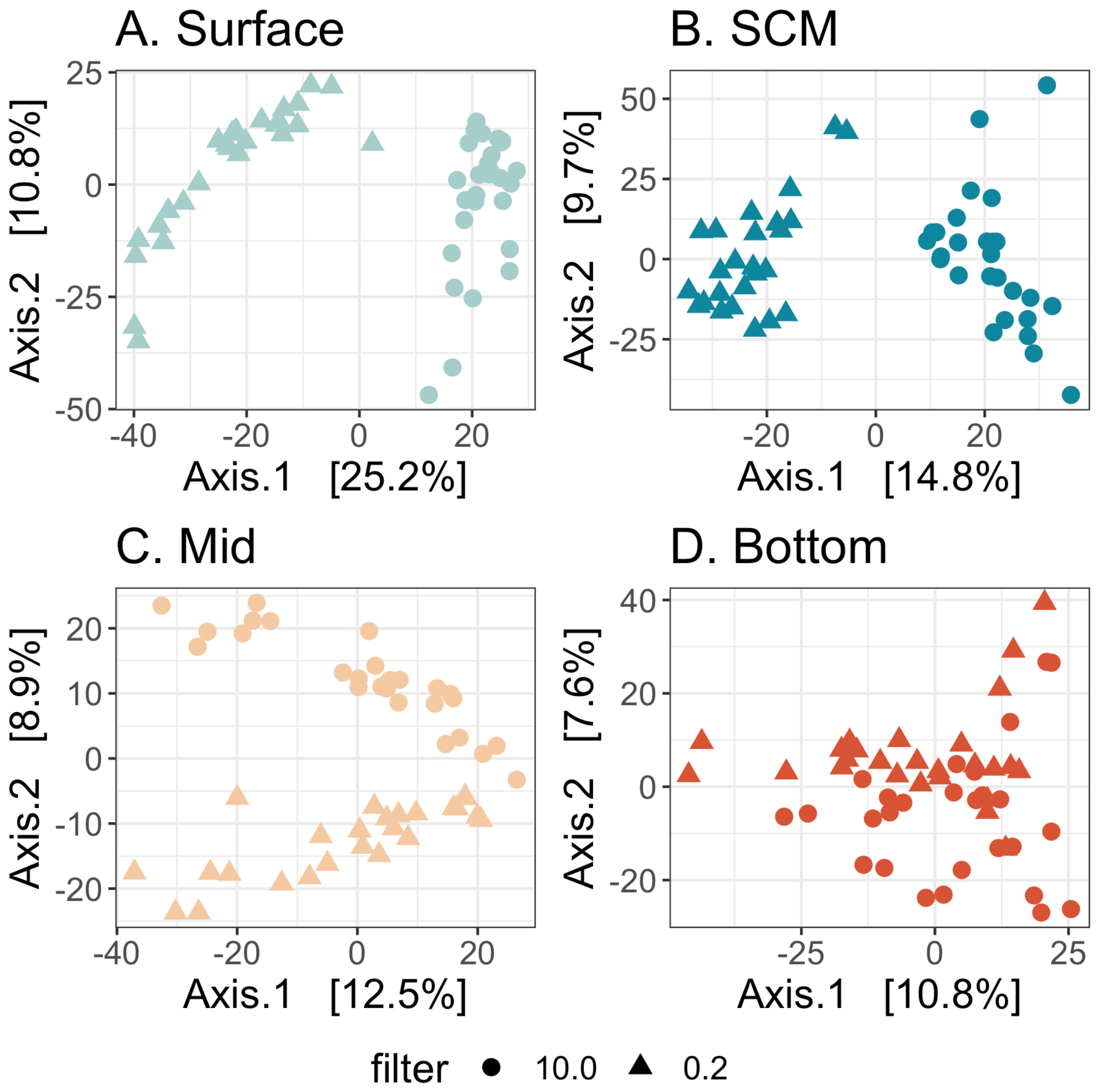


**Supplemental Figure 5. Principal coordinates analysis of Aitchison distances between size-fractionated protist communities from four depth layers in the Okinawa Trough.** PCoA were plotted by depth layer to better resolve the effect of filter pore-size on community composition. Point shape reflects the filter pore-size used to collect samples in μm; circles represent 10.0 μm filter pore-size and triangles represent 0.2 μm filter pore-size. Samples collected from the surface (A), SCM (B) clustered by filter pore-size on the primary axis samples collected from mid (C) and bottom (D) waters clustered by filter-pore size on the secondary axis. PERMANOVA results by filter pore-size were not significant for any of the depths.

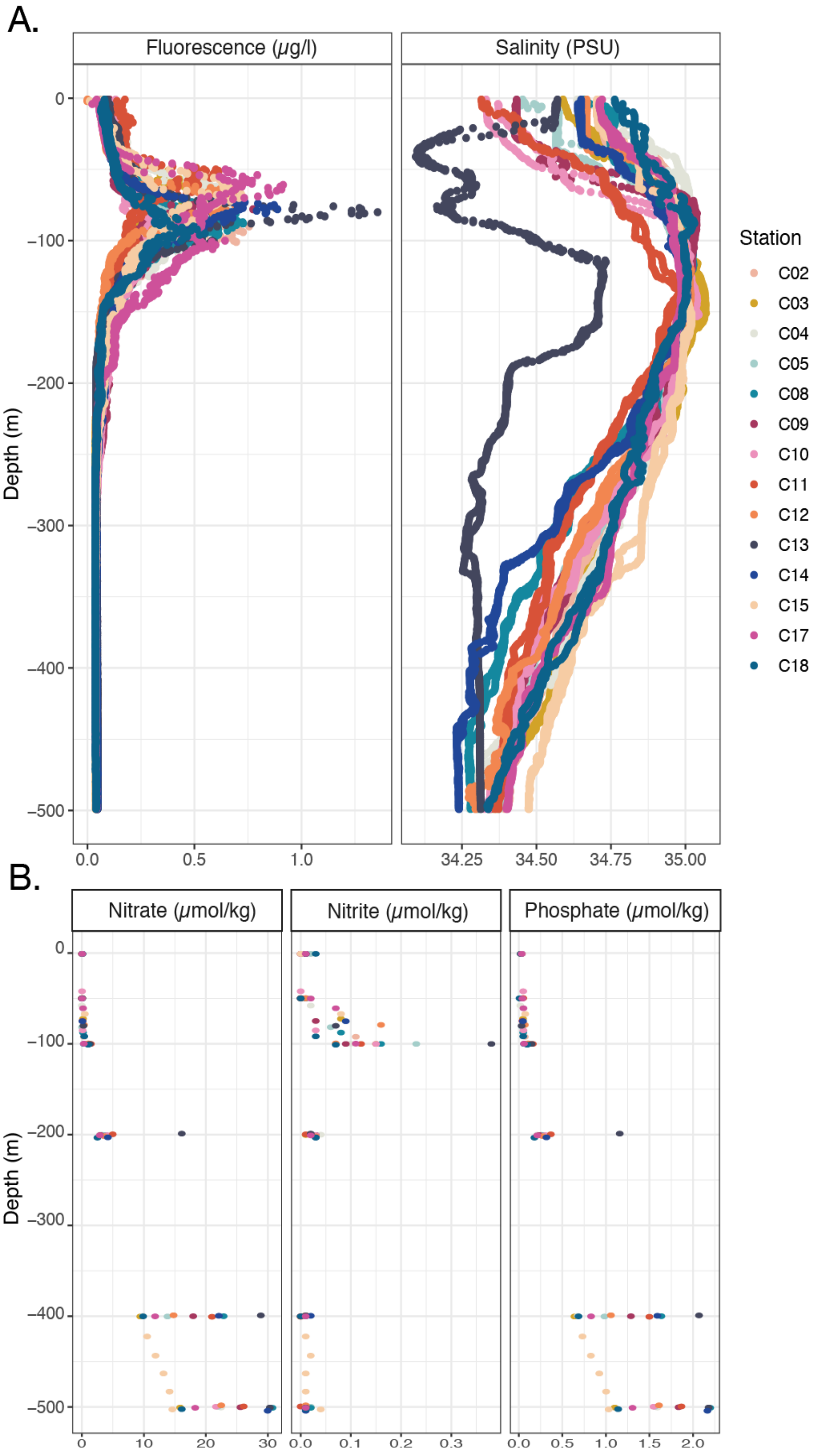


**Supplementary Figure 6. Depth Profiles for fluorescence (µg/l), salinity (PSU) and nutrient concentrations (µmol/kg) in the top 500 m for all stations.** Salinity and fluorescence (A) were measured directly by CTD sensors. Nitrate, nitrite, and phosphate (B) were analyzed for discrete water samples collected by Niskin bottle. Marine Works Japan Ocean Chemistry Analysis Section completed chemical analyses. Station 13 has the highest fluorescence values and lowest salinity at the SCM. Station 13 additionally has the highest nitrite concentration just below the SCM (-100 m) and high nitrate and phosphate concentrations below the SCM (-200 m).
